# Integrated Multi-Omics Mapping of Mitochondrial Dysfunction and Substrate Preference in Barth Syndrome Cardiac Tissue

**DOI:** 10.1101/2025.04.17.649294

**Authors:** Bauke V. Schomakers, Adriana S. Passadouro, Maria M. Trętowicz, Pelle J. Simpson, Yorrick R.J. Jaspers, Michel van Weeghel, Iman Man Hu, Cathelijne M. E. Lamboo, Denise Cloutier, Barry J. Byrne, Jan Bert van Klinken, Paul M. L. Janssen, Sander R. Piersma, Connie R. Jimenez, Frédéric M. Vaz, Gajja S. Salomons, Jolanda van der Velden, Riekelt H. Houtkooper, Signe Mosegaard

**Affiliations:** Laboratory Genetic Metabolic Diseases, Amsterdam UMC, University of Amsterdam, Amsterdam, The Netherlands; Core Facility Metabolomics, Amsterdam UMC, University of Amsterdam, Amsterdam, The Netherlands; Amsterdam Cardiovascular Sciences, Amsterdam, The Netherlands; Amsterdam Gastroenterology, Endocrinology and Metabolism, Amsterdam, The Netherlands; Department of Physiology, Amsterdam UMC, Vrije Universiteit Amsterdam, Amsterdam, The Netherlands; Enveda, Boulder, Colorado; Department of Pediatrics in the College of Medicine, University of Florida, Gainesville, FL; Laboratory for General Clinical Chemistry, Amsterdam UMC, University of Amsterdam, Amsterdam, The Netherlands; Department of Physiology and Cell Biology, College of Medicine, The Ohio State University, Columbus, Ohio, USA; OncoProteomics Laboratory, Amsterdam UMC, Location VUmc, Medical Oncology, Amsterdam, The Netherlands; Emma Center for Personalized Medicine, Amsterdam, The Netherlands

**Keywords:** Barth Syndrome, Metabolic Diseases, OMICS, Cardiomyopathy

## Abstract

Barth syndrome (BTHS) is a rare X-linked recessively inherited disorder caused by variants in the TAFAZZIN gene, leading to impaired conversion of monolysocardiolipin (MLCL) into mature cardiolipin (CL). Accumulation of MLCL and CL deficiency are diagnostic markers for BTHS. Clinically, BTHS includes cardiomyopathy, skeletal myopathy, neutropenia, and growth delays. Severely affected patients may require early cardiac transplants due to unpredictable cardiac phenotypes. The pathophysiological mechanisms of BTHS are poorly understood, and treatments remain symptomatic.

This study analyzed heart samples from five pediatric male BTHS patients (5 months-15 years) and compared them to tissues from 24 non-failing donors (19-71 years) using an integrated omics method combining metabolomics, lipidomics, and proteomics. The analysis confirmed changes in diagnostic markers (CL and MLCL), severe mitochondrial alterations, metabolic shifts, and elevated heart-failure markers. It also revealed significant interindividual differences among BTHS patients. With this study describe a powerful analytical tool for the in-depth analysis of metabolic disorders and a solid foundation for the understanding of BTHS disease phenotypes in cardiac tissues.

## Introduction

Barth Syndrome (MIM: 302060) is a rare X-linked recessively inherited mitochondrial disorder caused by pathogenic variants in *TAFAZZIN*^1,2^. The early onset of this disorder includes cardiomyopathy (73% in prenatal period), skeletal muscle myopathies, neutropenia and growth and developmental delays^3,4^.

*TAFAZZIN* encodes the Tafazzin protein, a mitochondrial transacylase that is crucial during the cardiolipin (CL) maturation process. CL is a phospholipid of the mitochondrial inner membrane (IM) essential for this membrane’s architecture and mitochondrial structure^5^. After synthesis, CL undergoes remodeling to establish a unique fatty acid composition that is tightly connected to its function in mitochondria. During this process, nascent CL is deacylated to monolysocardiolipin (MLCL) which is then re-acylated by Tafazzin to generate mature CL^6^. As CL contains four fatty acids and these can all be distinct, there can be many different CL species per tissue or cell type. In the mammalian heart, there is a specific species composition where linoleic acid (18:2) is the predominant fatty acid^6^. Tafazzin dysfunction leads to the MLCL accumulation and CL depletion since the remodeling process is impaired, making an elevated MLCL/CL ratio a highly specific BTHS diagnostic biomarker^7^.

Cardiomyopathy is the most common clinical presentation in infants affected with BTHS and, in the early childhood period, cardiac transplant might be required^3,8^. This could be the case for 12-26% of BTHS individuals as estimated previously^3,9^. The cardiac presentations can be diverse and unpredictable involving ventricular arrhythmias (10-44%), dilated cardiomyopathy (DCM) (96%), hypertrophic cardiomyopathy (HCM) (3%), restrictive cardiomyopathy, left ventricular noncompaction (19-53%), endocardial fibroelastosis and sudden cardiac death^3,4,10,11^. Standard heart failure (HF) medications have proven to stabilize BTHS patients^3,4,12^. However, after years of cardiac stability, mechanical support and cardiac transplant may still be required despite the risk of a major operation in patients suffering from neutropenia^12–14^. The lack of treatment options and the severity of the phenotype so early in life stimulated scientific research into new possible therapeutic avenues, focusing traditionally on animal models.

Previously, male *TAFAZZIN*-knockdown (KD) mouse models in C57BL/6 genetic background were established and the proteome of cardiac mitochondria isolated from ventricular myocardium was obtained. Proteins related to oxidative phosphorylation (OXPHOS) and ubiquinone biosynthesis were reduced in cardiac mitochondrial^15^. The electron transport chain (ETC) supercomplexes were destabilized and, consequently, the interactions between these supercomplexes and the fatty acid oxidation (FAO) enzymes were abnormal^15^. When analysing the lipid profile of *TAFAZZIN*-KD mice, a sharp decrease in linoleic acid containing molecular species was identified^16^. Tafazzin knock-down was ultimately associated with alterations in choline and ethanolamine glycerophospholipids and triggered an alteration of myocardial substrate utilization from fatty acids and glucose to amino acid oxidation^16^.

While these animal models have provided novel insights, the translational gap to humans is considerable^17^. Human studies have mainly focused on sample types not directly involved in cardiac function, such as lymphoblastoid cells^18^, plasma^19^ and fibroblasts^20^. Previously, LC-MS/MS was used to establish the lipid profile of lymphoblastoid cell pellets from five BTHS patients^18^. Dilysocardiolipin (DLCL), MLCL and CL levels were aberrant, as well as several other lipid species. In a larger cohort, plasma from 23 BTHS patients was analyzed using NMR metabolomics and targeted LC-MS metabolomics^19^. Clear differences between BTHS and control samples included acylcarnitines, amino acids, biogenic amines, and glycerophospholipids. Mitochondria were isolated from primary skin fibroblasts of BTHS patients and used for complexome profiling and fluxomics^20^. Partial destabilization of the supercomplexes was detected along with a substrate preference shift towards glutamine. Importantly, it was suggested that further studies were required in order to assess the functional effects these observations could have on high energy tissues, considering that BTHS fibroblasts seemed minimally affected^20^.

In the present study, we have used an integrative omics approach on left ventricular heart tissue samples from five individuals with BTHS for an in-depth analysis of the BTHS cardiac phenotype. The same five cardiac samples were previously analyzed using LC-MS/MS to investigate the BTHS cardiac proteome between BTHS cardiomyopathy and idiopathic DCM in children, identifying that long-chain FAO (lcFAO) was impaired and glucose uptake and utilization was increased^21^. To expand on these findings and map the broad cellular consequences of BTHS, we employed a state-of-the-art multi-omics strategy involving (a) semi-targeted metabolomics, (b) lipidomics of complex lipids, and (c) analytical flow DIA proteomics. All omics methods were performed using a single and unified sample preparation, minimizing sampling variability.

## Results

### Integrative OMICs show phenotypic differences in human BTHS heart samples

In this study, the metabolic, lipidomic and proteomic profiles of five left ventricular heart samples from BTHS individuals and 24 hearts from non-failing donors were studied (Supplemental Table 1). The BTHS donors were male, aged between 5 months and 15 years old. Two samples were obtained during autopsy and three at heart transplantation. The Principal Component Analyses (PCA) of metabolomics (123 metabolites, Supplemental Table 2), lipidomics (1899 lipids, Supplemental Table 3) and proteomics (1445 proteins, after QC, Supplemental Table 4) results (Figure 1A-C) show BTHS triplicates (separately prepared samples from the same donors) individually, allowing for an assessment of their experimental variance. The PCA plots show a clear separation of BTHS individuals from the non-failing donors, except for transplantation samples in metabolomics. In each of the omics methods, hearts from the two patients obtained at autopsy showed wider separation from non-failing donors than those that were obtained after transplantation (Figure 1A-C). These findings highlight not only the clear distinction between non-failing donors and BTHS individuals but also the complexity and heterogeneity of BTHS, with various cellular processes being differentially affected in patients.

**Figure 1.**
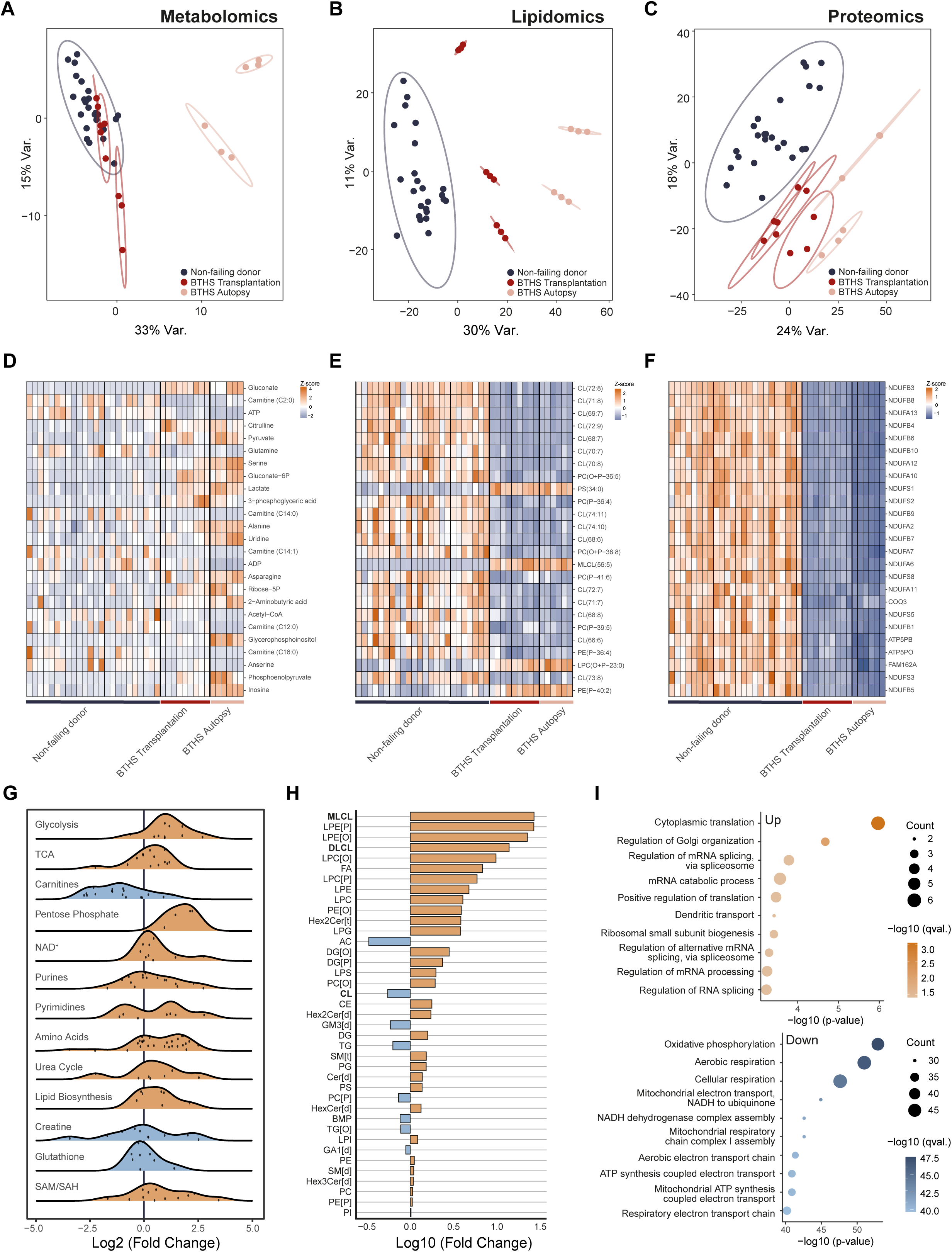
Overview of phenotypic differences between BTHS (n=5) and non-failing donors (n=24) detected in metabolomics, lipidomics and proteomics. **(A-C)** Principal Component Analysis (PCA) for metabolomic (A), lipidomic (B) and proteomic (C) profiles, showing three technical BTHS replicates. Non-failing donors show distinct separation from all BTHS tissues, except for BTHS samples obtained during transplantation in metabolomics. BTHS tissues obtained at autopsy show a larger separation from non-failing donors than those obtained during transplantation in all analyses. **(D-F)** Heatmaps representing z-scores of the top 25 (ranked on P-value) metabolites (D), lipids (E) and proteins (F) in cardiac samples of BTHS individuals compared to non-failing donors. **(G)** Ridge plot of the changes in (polar) metabolic pathways shows pleiotropic differences between tissues of BTHS patients and non-failing donors, especially in energy related pathways. **(H)** Lipid class totals across cardiac samples from BTHS individuals and non-failing donors, highlighting lipids related to the Tafazzin deficiency such as MLCL, DLCL and CL. **(I)** Bubble plot of enriched gene ontology (GO) biological processes for upregulated (top panel) and downregulated (bottom panel) proteins in BTHS cardiac tissue indicate that mitochondrial respiration is significantly decreased.

Next, we studied the top 25 metabolites, lipids and proteins sorted on *P*-Value (Figure 1D-F). Metabolite ranking is based on a comparison between non-failing donors and the fifteen BTHS samples (five individuals in biological triplicates) as a single group. Polar metabolites show homogeneity within patient replicates (Figure 1D). However, their intensities are diverse between each individual, and several metabolites are significantly elevated in the samples collected at autopsy. In lipidomics, (monolyso-) cardiolipins dominate the list of significantly regulated lipids and show high uniformity in all BTHS samples (Figure 1E). The PCA analysis demonstrate a high degree of separation even within the BTHS group, which hints at deeper effects on the lipidome not represented in this heatmap, as that shows a more uniform picture. Proteomics reveals a very clear distinction in protein abundances between controls and BTHS individuals, with the top significant changes being dominated by proteins from the NDUF family which belong to respiratory chain complex I (Figure 1F). Finally, to illustrate the extent of changes within the metabolome, lipidome and proteome in BTHS heart tissue compared to non-failure control cardiac tissue, we compared both groups using metabolic pathways, lipid classes and GO-terms as classifying features respectively (Figure 1G-I). From the metabolomics data, differences are linked to pathways related to energy, such as glycolysis, short chain carnitines and nucleotides (Figure 1G), which corroborates the results in the top 25 altered metabolites. When surveying all lipids classes, CL and MLCL, directly related to the Tafazzin deficiency that causes BTHS are indeed highly affected (Figure 1H). This overview reveals the depth of the alterations in the lipidome. Strikingly, many lysolipid-species of glycerophospholipids such as phosphatidylethanolamine (PE) and phosphatidylcholine (PC), including their alkyl and plasmalogen variants, are highly increased in abundance. While previous studies in the TAFAZZIN knockdown mouse heart showed unchanged PE[P] and reduced PC[P], our analysis in human heart tissue confirms these findings but additionally reveals elevated lyso-ether lipids (LPE[O], LPE[P], and LPC counterparts), which were not measured in the mouse model^22,23^. In proteomics, GO-term enrichment analysis highlights the expected depletion in proteins related to mitochondrial respiration (Figure 1I). Notably, there are far fewer proteins increased than decreased, and to a much lesser degree, with no clear trends. After determining that the replicates show only minor variance, their average abundances for metabolites, lipids or proteins were used for further specific analysis and statistics.

### Confirmation of diagnostic markers CL, MLCL and DLCL in BTHS cardiac tissue

BTHS is caused by a deficiency in the cardiolipin remodelling enzyme Tafazzin (Figure 2A). Clinical diagnosis of BTHS is therefore often initiated by determining the ratio of CL(72:8) and MLCL(50:2) in blood plasma or blood spots, as mature CL is depleted and the MLCL precursor is highly accumulated^7^. This distinctive profile is also clearly seen in cardiac tissue (Figure 2B). A broader analysis reveals that the CL lipid cluster shows major disruption (Figure 2C), with a marked decrease in mature CL species, and a corresponding increase in immature CL species. Statistical analysis of individual MLCL and DLCL species is not possible, as these are below the limits of detection in healthy individuals. However, total MLCL and DLCL levels clearly illustrate their accumulation in BTHS affected individuals (Figure 2D). Notably, for both the clinically validated biomarkers and the total levels of MLCL and DLCL, there are no obvious correlation between their levels and whether the heart sample was obtained at transplantation or post-mortem.

**Figure 2.**
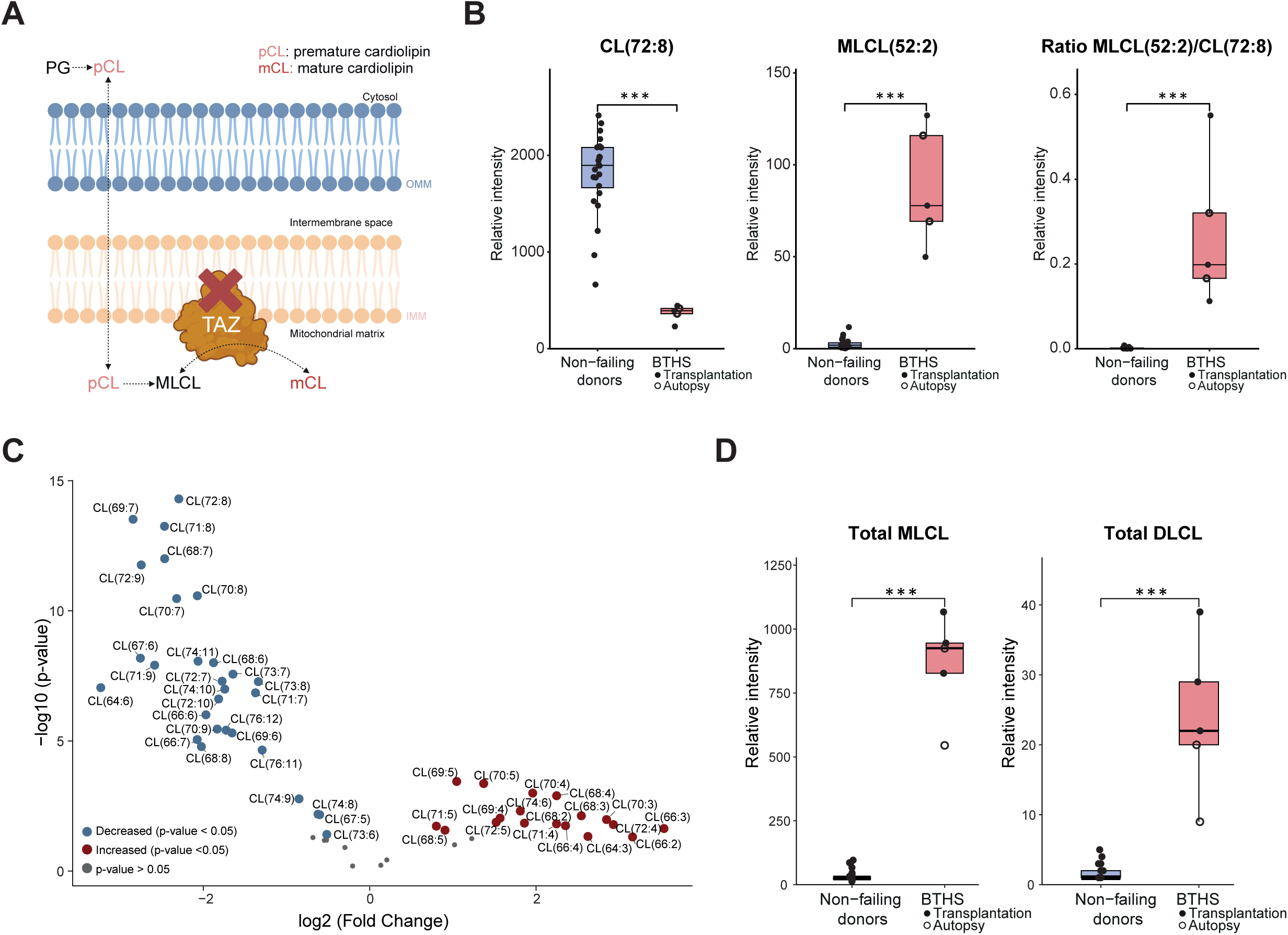
Diagnostic markers CL, MLCL and DLCL in BTHS cardiac tissue. **(A)** Overview of cardiolipin biosynthesis and remodeling via the Tafazzin enzyme, which is dysfunctional in BTHS. Created with Biorender.com. **(B)** Relative intensity of CL(72:8), MLCL(50:2) and the ratio between MLCL(50:2) and CL(72:8) in both BTHS and non-failing donors. **(C)** Volcano plot of CL species, depicting depleted CL species (blue) and accumulated CL species (red) in cardiac tissue of BTHS individuals compared to non-failing donors**. (D)** Relative intensity of the total MLCL and DLCL classes in BTHS and non-failing donors.

### Integrative omics reveals mitochondrial dysfunction in cardiac tissue from BTHS affected individuals

Mitochondrial dysfunction is a hallmark of BTHS and previous reports have shown structural abnormalities^24,25^ and OXPHOS dysfunction that result in energy depletion^12,26^ (Figure 3A). As cardiolipin is a fundamental part of the mitochondrial membrane, we analysed lipid classes associated with the mitochondrial membrane from our lipidomics dataset^27^. Interestingly, only cardiolipin is strongly altered (Figure 3B). As CL make up a significant part of the mitochondrial membrane, we analysed a subset of mitochondrial proteins according to the Mitocarta 3.0 database^28^. This comparison highlights striking alterations in cardiac mitochondrial proteins from BTHS individuals (Figure 3C). Cardiolipins play a critical role in stabilizing electron transport complexes^5^ (Figure 3A), and our GO-term analysis identified a significant reduction of OXPHOS proteins (Figure 2I). Consistently, proteins associated with mitochondrial OXPHOS complexes are markedly decreased (Figure 3D). In fact, a majority of proteins in complex I and V are reduced. Interestingly, all CoQ enzymes in the current dataset, involved in the production of CoQ10, an important electron carrier in the ETC, are significantly reduced (Figure 3E). However, CoQ10 itself is maintained at normal levels. Finally, a reduction in energy carriers such as adenosine triphosphate (ATP) shows a clear trend (Figure 3F). The same pattern is also observed in phosphocreatine, a phosphate storage for rapid ATP production, with two of the critical enzymes related to this phosphate exchange significantly decreased (Figure 3G).

**Figure 3.**
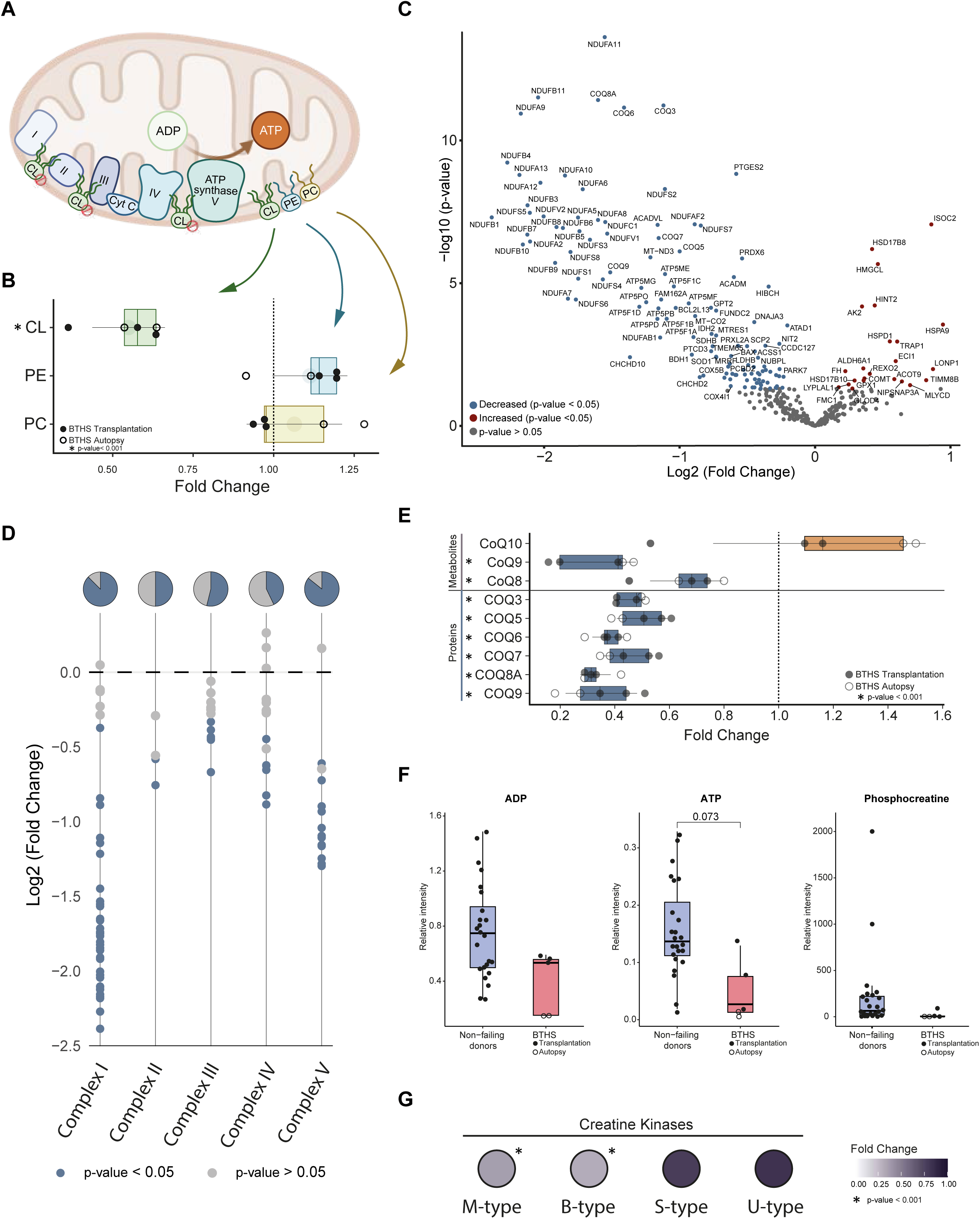
Mitochondrial dysfunction in cardiac tissue from BTHS individuals. **(A)** Schematic overview of the mitochondria, highlighting the function of CL as a mitochondrial membrane lipid. In BTHS, CL depletion contributes to the destabilization of the ETC complexes which impairs ATP production via OXPHOS. Created with Biorender.com. **(B)** Changes in abundance of main mitochondrial membrane lipids in cardiac tissue of BTHS individuals compared to non-failing donors. **(C)** Volcano plot of mitochondrial proteins (according to MitoCarta 3.0), depicting depleted- (blue) and accumulated proteins (red) in cardiac tissue of BTHS individuals compared to non-failing donors. **(D)** Changes in electron transport chain proteins show major dysfunction, and a majority of decreased proteins in Complex I and Complex V. The pie charts represent the proportion of decreased proteins in each complex. **(E)** Changes in coenzyme Q related metabolites and proteins, associated with the electron transport chain, in cardiac tissue of BTHS individuals compared to non-failing donors. **(F)** Relative intensity of ADP, ATP and phosphocreatine in both BTHS and non-failing donors. **(G)** Changes in abundance of creatine kinase isoforms of M-type, B-type, S-type, U-type in BTHS cardiac tissues where colour intensity corresponds to the magnitude of change in comparison to non-failing donors.

### BTHS cardiac tissue shows a metabolic substrate shift

In a healthy heart, myocardial metabolism primarily relies on FAO to meet its high energy demand^29^. As we observed severe depletion of mitochondrial proteins especially, as well as ATP and other energy related metabolites, we studied the energy metabolism of BTHS tissues. Combining data from all three omics, we provide a comprehensive overview of cellular energy metabolism (Figure 4), including glycolysis (Figure 4A), the TCA cycle (Figure 4B) and mitochondrial β-oxidation (Figure 4C). We observed that acetyl-CoA, the main substrate of the TCA cycle, is significantly depleted in BTHS cardiac tissue. Acetyl-CoA can be produced through various pathways, with the predominant sources being FAO and glycolysis. In line with a metabolic shift away from FAO, we observe a major and largely significant decrease of carnitines, independent of chain length and saturation. Several proteins related to mitochondrial β-oxidation show a decreased abundance. Very long-chain acyl-CoA dehydrogenase (VLCAD), responsible for the oxidation of long chain fatty acids within mitochondria, showed the greatest reduction, followed by medium-chain acyl-CoA dehydrogenase (MCAD) whereas short-chain acyl-CoA dehydrogenase (SCAD) shows no changes. Strikingly, individuals with reduced levels of these proteins displayed an approximately two-fold increase in the trifunctional protein HADHA, a major component of mitochondrial β -oxidation, whereas no such increase was observed in other BTHS individuals. Along with this apparent general decrease in FAO in BTHS, there is a complementary increase in the glycolytic end-products pyruvate and lactate, as well as alanine. Interestingly, glycolysis enzyme abundance seems to show pleiotropic dysfunction. Hexokinase 1 and the Pyruvate dehydrogenase complex show a trend towards upregulation, while four of the intermediate enzymes are significantly downregulated. Finally, TCA cycle metabolites appeared unaffected, while related enzymes show a trend of downregulation. Although considerable inter-individual variation was observed, the results suggest an altered substrate preference from FAO to glycolysis in BTHS heart tissue.

**Figure 4.**
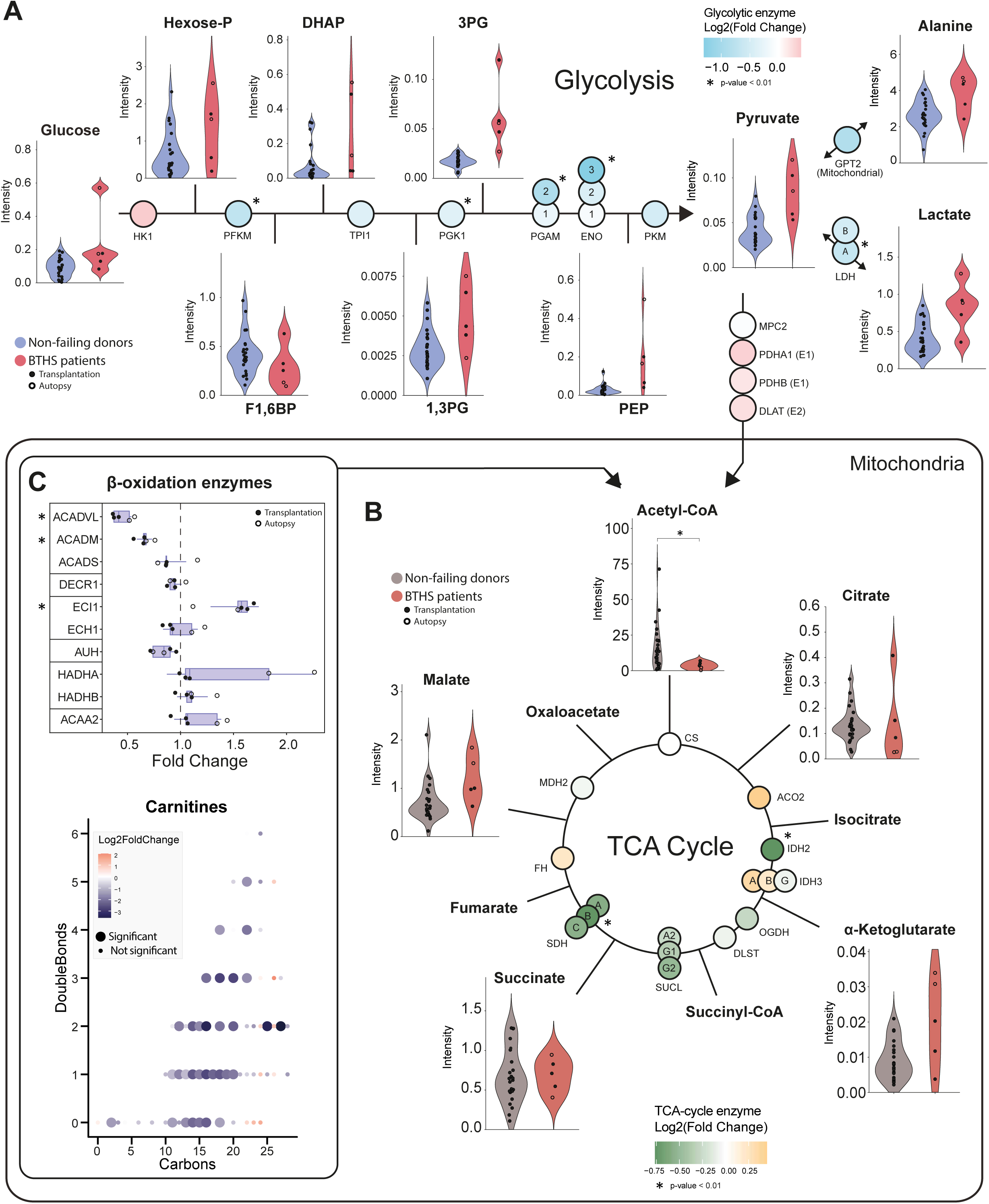
Comprehensive overview of energy metabolism in BTHS cardiac tissue, integrating metabolomics, lipidomics and proteomics data. The interconnections between **(A)** glycolysis, **(B)** the TCA cycle and **(C)** β -oxidation are shown. For glycolysis and the TCA cycle, relative metabolite intensity is represented in the violin plots comparing the non-failing donors and the BTHS group. Changes in the abundance of enzymes are represented using a colour gradient in the circles between metabolites. β-Oxidation enzymes are shown as horizontal box-plots, and an overview is provided of changes in carnitine species. Abbreviations: F1,6BP; Fructose-1,6-biphosphate, DHAP; Dihydroxyacetone phosphate, 1,3PG; 1,3-phosphoglycerate, 3PG; 3-phosphoglycerate, PEP; Phosphoenolpyruvate.

### Cardiac remodelling in BTHS hearts

BTHS patients often present with diminished cardiac performance, though individual phenotypes are diverse^3,6^. To investigate whether BTHS hearts undergo structural remodelling, we analysed actin-myosin cytoskeleton proteins (Figure 5A) and compared their abundance between BTHS tissues obtained at autopsy and during transplantation (Figure 5B). BTHS hearts exhibit numerous altered proteins associated with cardiac structure as well as electrical and mechanical function. In line with this, metabolic changes associated with remodelling and heart failure were observed in BTHS hearts. Alterations in branched-chain amino acid (BCAA) metabolism have been observed in HF models^30,31^. BCAA were markedly increased in the two deceased individuals, compared to both controls and samples obtained at transplantation (Figure 5C). Enzymes related to the metabolism of BCAAs also exhibit a noticeable difference between deceased and other individuals (Figure 5D). Furthermore, lipidomics revealed substantial accumulation of heart failure associated lipids, mostly in deceased individuals, such as sphingosines (Figure 5E), ceramides (Figure 5F) and fatty acids (Figure 5G). These types of lipids can contribute to lipotoxicity when accumulated in cardiac tissue^32–35^. In BTHS, total sphingosine and ceramide levels were doubled in individuals with failing hearts compared to non-failing controls. Interestingly, these levels were within the normal range for BTHS patients without heart failure. Fatty acids were increased in all BTHS tissues but were significantly increased in deceased individuals. When looking at the composition of these lipids, mostly the shorter sphingosines appear affected, while ceramides and fatty acids are more uniformly increased.

**Figure 5.**
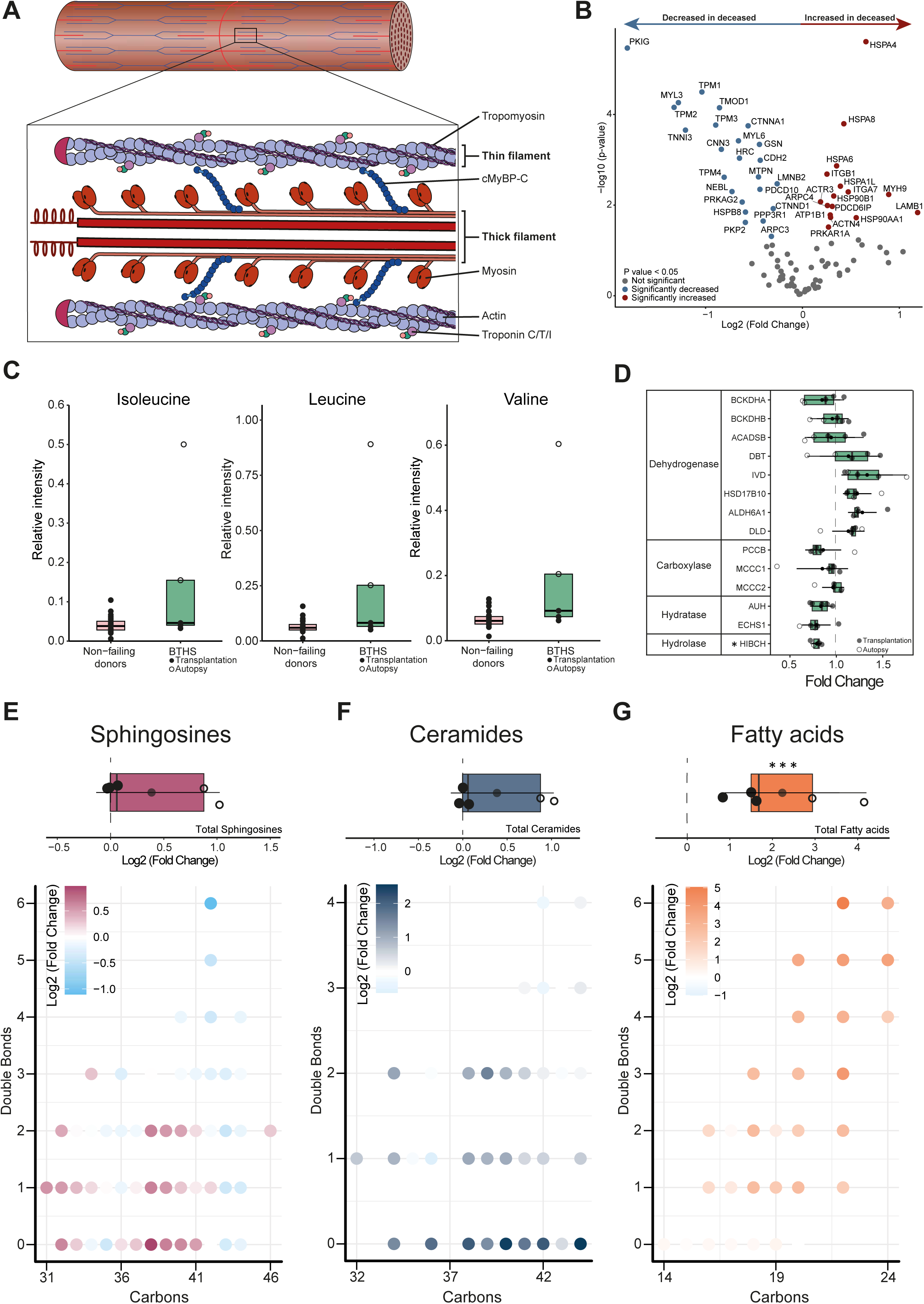
Cardiac remodelling in BTHS cardiac tissue. **(A)** Structure of cardiac actin-myosin cytoskeleton. **(B)** Volcano plot of cardiac structural proteins depicting depleted proteins (blue) and accumulated proteins (red) in cardiac tissue of BTHS individuals obtained at autopsy (n=2) and transplantation (n=3). **(C)** Relative intensity of isoleucine, leucine and valine in both BTHS and non-failing donors groups. **(D)** Changes in enzymes involved in branched-chain amino acid metabolism in cardiac tissue of BTHS individuals when compared to non-failing donors. **(E)** At the top, changes in total sphingosines in cardiac tissue of BTHS individuals when compared to non-failing donors. At the bottom, an overview of changes in individual sphingosine species when comparing the same groups. This is repeated for **(F)** Ceramides and **(G)** Fatty acids.

## Discussion

The aim of this study was to analyze the metabolome, lipidome and proteome of BTHS cardiac tissue and gain insights into the pathological mechanisms behind BTHS cardiomyopathy in human hearts. Our combined triple-omics platform is a powerful analytical tool since the direct comparison and integration of the multi-omics results is greatly aided by the fact that all omics analyses were performed on a single sample of heart tissue. This approach allowed us to achieve a comprehensive molecular and metabolic characterization of BTHS cardiac tissue. Corroborating earlier findings, our method detected the effects of the primary defect, accumulation of MLCL (and DLCL), CL depletion^36,37^.

Observed across all three datasets, BTHS heart samples showed clear signs of mitochondrial dysfunction and energy depletion. The proteome of BTHS heart samples showed the NDUF protein family significantly decreased. These proteins are subunits of respiratory complex I, and their depletion is a known hallmark in BTHS, previously reported in lymphoblasts^38^ and fibroblasts^20^ from BTHS individuals. Additionally, proteins in the complexes I to V were severely decreased. This was expected since the absence of mature CL, or accumulation of MLCL, can impact mitochondrial inner membrane architecture and also the integrity and enzymatic activity of the ETC supercomplexes that generate ATP through OXPHOS^15,39,40^. Changes in the CL pool have also been causally linked to structural abnormalities of the ETC supercomplexes in heart failure^41,42^. In the metabolic profile, we also identified the downstream effects with ADP, ATP and creatine phosphate showing noticeable, though not significant, decreases in BTHS. The latter is known to be an energy buffer to ensure myofilament contractility in heart tissue and the autopsy samples from BTHS individuals seem to be most affected by this apparent inability to maintain energy homeostasis^43^.

Considering the significant metabolic flexibility the human heart has to properly adapt to substrate availability and uptake, we investigated sugar and fat metabolism^32,44^. The heart relies mainly on fatty acids for ATP production (60-90%)^29^, and a shift from FAO towards glucose oxidation can represent a maladaptive compensatory mechanism^45^. In this study, BTHS cardiac tissues showed profound and pleiotropic effects associated with mitochondrial function and energy production. We observed a reduction in FAO related metabolites and enzymes in BTHS heart samples. This mechanism was described earlier in a BTHS mouse model^46^, identified in cardiomyocytes derived from BTHS patient-induced pluripotent stem cells (iPSC)^46^ and suggested to be directly associated with worse cardiac function in young adults with BTHS^47^.

Metabolic remodelling and structural remodelling are tightly connected in heart tissue^45^, which was also observed in this study. The actin-myosin cytoskeleton showed signs of structural remodelling when comparing heart tissue samples obtained at autopsy to those obtained at transplantation. These alterations were in line with findings from BTHS patient-derived cellular models, where mitochondrial abnormalities have been reported to cause sarcomere disarray and subsequent defective contractility^48^.

We observed alterations in amino acid metabolism in the BTHS heart samples. However, these abnormalities were not consistent for all BTHS samples, as autopsy-derived and transplantation-derived samples clustered into two distinct groups. Notably, irregular myocardial amino acid metabolism has been reported in non-BTHS models of heart failure^30,31^. Furthermore, BCAA have been reported to accumulate in human DCM heart tissue^49^. Consistent with this our analysis revealed that BCAA levels were most significantly altered in tissues from deceased patients. This trend was mirrored by similar alterations in enzymes levels involved in BCAA metabolism.

A variety of metabolites and lipids associated with heart remodelling and failure were found to accumulate in BTHS heart samples, corroborating earlier findings in heart failure studies^50,51^. Sphingolipids and ceramides have been linked to cardiac lipotoxicity as they alter cellular membrane organization, inducing cardiomyocyte apoptosis^32,33^. Accumulated fatty acids contribute to lipotoxicity increasing ROS production that causes endoplasmic reticulum (ER) stress^34,35^. However, our integrated omics data also highlights the heterogeneity within the BTHS group. This accumulation of toxic lipids was especially clear in BTHS hearts collected at autopsy when compared to BTHS heart samples collected at transplantation, providing a possible lead for biomarkers related to disease severity.

A limitation of this study is the unavailability of age- and sex-matched heart tissue donor material. Due to the exceptionally young age at which BTHS heart failure occurs, matching cardiac material from healthy donors is exceedingly rare. Thus, our control group consists of heart tissue from a wide variety of phenotypes (male/ females and three different age groups). Despite this variability, we still identified consistent differences between both groups, highlighting a distinct BTHS phenotype. However, due to the diversity of our control group, more general markers of cardiac and mitochondrial dysfunction might not be identified to their full extent in the current comparisons. For instance, this could support the observed lack of glutathione depletion or clear shift towards glycolysis in BTHS hearts when compared to non-failing donors, despite severe mitochondrial dysfunction and likely lipotoxicity in BTHS hearts. Clinical heterogeneity is a known feature inherent in BTHS^3^ and the present study includes a relatively small sample size due to the inherent rarity of BTHS heart tissue samples. Two of these samples were obtained at autopsy and three at transplantation, thus likely under different circumstances. Unfortunately, patient information is limited and information such as *TAFAZZIN* genotype, medication and cardiac phenotype at time of collection are unknown^21^. Despite these limitations, our approach provides an unprecedented amount of high-quality data on the BTHS cardiac tissue, encompassing broad swaths of the heart tissues’ biological organization. The findings are consistent with prior studies on BTHS model systems and collectively reveal the extensive metabolic, lipidomic and proteomic alterations in BTHS heart tissues, serving as a robust foundation for future explorations into targeted treatment development.

In conclusion, we analysed the myocardial metabolic, lipidomic and proteomic profile of a single cardiac biopsy of individuals affected with BTHS using a unified extraction providing high quality data, confirming known BTHS aberrations in heart tissue, while also providing potentially new areas of interest.

## Methods

### Patient samples

Five left ventricular heart samples from individuals affected by BTHS cardiomyopathy were used in this study^21^. In Supplemental Table 1, the information about each sample is outlined. These samples were provided by the Barth Syndrome Foundation DNA and Tissue Bank. The samples from BTHS individuals were compared to a group of samples from non-failing donors (female/male; 18->55 years; Caucasian/African-American) provided by Ohio State University. BTHS triplicates, prepared as separate samples from the same donor tissue, were created to allow for the assessment of both biological and technical variance.

### Multi-omics sample preparation

Metabolomics, lipidomics and proteomics were performed after a unified extraction method, combining several established methods^52–55^. In a 2 mL tube, containing approximately 3 mg of freeze-dried heart tissue, the following amounts of internal standard dissolved in water were added to each sample for metabolomics: adenosine-^15^N_5_-monophosphate (5 nmol), adenosine-^15^N_5_-triphosphate (5 nmol), D_4_-alanine (0.5 nmol), D_7_-arginine (0.5 nmol), D_3_-aspartic acid (0.5 nmol), D_3_-carnitine (0.5 nmol), D_4_-citric acid (0.5 nmol), ^13^C_1_-citrulline (0.5 nmol), ^13^C_6_-fructose-1,6-diphosphate (1 nmol), ^13^C_2_-glycine (5 nmol), guanosine-^15^N_5_-monophosphate (5 nmol), guanosine-^15^N_5_-triphosphate (5 nmol), ^13^C_6_-glucose (10 nmol), ^13^C_6_-glucose-6-phosphate (1 nmol), D_3_-glutamic acid (0.5 nmol), D_5_-glutamine (0.5 nmol), ^13^C_6_-isoleucine (0.5 nmol), D_3_-lactic acid (1 nmol), D_3_-leucine (0.5 nmol), D_4_-lysine (0.5 nmol), D_3_-methionine (0.5 nmol), D_6_-ornithine (0.5 nmol), D_5_-phenylalanine (0.5 nmol), D_7_-proline (0.5 nmol), ^13^C_3_-pyruvate (0.5 nmol), D_3_-serine (0.5 nmol), D_6_-succinic acid (0.5 nmol), D_4_-thymine (1 nmol), D_5_-tryptophan (0.5 nmol), D_4_-tyrosine (0.5 nmol), D_8_-valine (0.5 nmol).

In the same 2 mL tube, the following amounts of internal standards dissolved in 1:1 (v/v) methanol:chloroform were added for lipidomics: Bis(monoacylglycero)phosphate BMP(14:0)2 (0.2 nmol), Ceramide-1-phosphate C1P(d18:1/12:0) (0.125 nmol), D_7_-Cholesteryl Ester CE(16:0) (2.5 nmol), Ceramide Cer(d18:1/12:0) (0.125 nmol), Ceramide Cer(d18:1/25:0) (0.125 nmol), Cardiolipin CL(14:0)4 (0.1 nmol), Diacylglycerol DAG(14:0)2 (0.5 nmol), Glucose Ceramide GlcCer(d18:1/12:0) (0.125 nmol), Lactose Ceramide LacCer(d18:1/12:0) (0.125 nmol), Lysophosphatidicacid LPA(14:0) (0.1 nmol), Lysophosphatidylcholine LPC(14:0) (0.5 nmol), Lysophosphatidylethanolamine LPE(14:0) (0.1 nmol), Lysophosphatidylglycerol LPG(14:0) (0.02 nmol), Phosphatidic acid PA(14:0)2 (0.5 nmol), Phosphatidylcholine PC(14:0)2 (2 nmol), Phosphatidylethanolamine PE(14:0)2 (0.5 nmol), Phosphatidylglycerol PG(14:0)2 (0.1 nmol), Phosphatidylinositol PI(8:0)2 (0.5 nmol), Phosphatidylserine PS(14:0)2 (5 nmol), Sphinganine 1-phosphate S1P(d17:0) (0.125 nmol), Sphinganine-1-phosphate S1P(d17:1) (0.125 nmol), Ceramide phosphocholines SM(d18:1/12:0) (2.125 nmol), Sphingosine SPH(d17:0) (0.125 nmol), Sphingosine SPH(d17:1) (0.125 nmol), Triacylglycerol TAG(14:0)2 (0.5 nmol). Subsequently, solvents were added to achieve a total volume of 500 µL water, 500 µL methanol. Muscle tissues were homogenized using a Qiagen TissueLyser II for 2 minutes at 30 times/s with a 5 mm Qiagen Stainless Steel Bead in each tube. Chloroform was added to each tube for a total chloroform volume of 1 mL and samples were thoroughly mixed, before centrifugation for 10 min at 14.000 rpm to facilitate layer separation.

### Metabolomics

The top layer, containing the polar phase, was transferred to a new 1.5 mL tube and dried using a vacuum concentrator at 60°C. Dried samples were reconstituted in 100 µL 6:4 (v/v) methanol:water. Metabolites were analysed using a Waters Acquity ultra-high performance liquid chromatography system coupled to a Bruker Impact II™ Ultra-High Resolution Qq-Time-Of-Flight mass spectrometer. Samples were kept at 12°C during analysis and 5 µL of each sample was injected. Chromatographic separation was achieved using a Merck Millipore SeQuant ZIC-cHILIC column (PEEK 100 x 2.1 mm, 3 µm particle size). Column temperature was held at 30°C. Mobile phase consisted of (A) 1:9 (v/v) acetonitrile:water and (B) 9:1 (v/v) acetonitrile:water, both containing 5 mmol/L ammonium acetate. Using a flow rate of 0.25 mL/min, the LC gradient consisted of: Dwell at 100% Solvent B, 0-2 min; Ramp to 54% Solvent B at 13.5 min; Ramp to 0% Solvent B at 13.51 min; Dwell at 0% Solvent B, 13.51-19 min; Ramp to 100% B at 19.01 min; Dwell at 100% Solvent B, 19.01-19.5 min. Column was equilibrated by increasing flow rate to 0.4 mL/min at 100% B for 19.5-21 min. MS data were acquired using negative and positive ionization in full scan mode over the range of m/z 50-1200. Data were analysed using Bruker TASQ software version 2.1.22.3. All reported metabolite intensities were normalized to freeze-dried tissue weight, as well as to internal standards with comparable retention times and response in the MS. Metabolite identification was based on a combination of accurate mass, (relative) retention times and fragmentation spectra, compared with the analysis of a library of standards.

### Lipidomics

The bottom layer, containing the apolar phase, was transferred to a new 1.5 mL tube and evaporated under nitrogen at 60°C. The residue was dissolved in 100 μL of 1:1 (v/v) methanol:chloroform. Lipids were analysed using a Thermo Scientific Ultimate 3000 binary HPLC coupled to a Q Exactive Plus Orbitrap mass spectrometer. For normal phase separation, 2 μL of each sample was injected onto a Phenomenex® LUNA silica, 250 * 2 mm, 5µm 100Å. Column temperature was held at 25°C. Mobile phase consisted of (A) 85:15 (v/v) methanol:water containing 0.0125% formic acid and 3.35 mmol/L ammonia and (B) 97:3 (v/v) chloroform:methanol containing 0.0125% formic acid. Using a flow rate of 0.3 mL/min, the LC gradient consisted of: Dwell at 10% A 0-1 min, ramp to 20% A at 4 min, ramp to 85% A at 12 min, ramp to 100% A at 12.1 min, dwell at 100% A 12.1-14 min, ramp to 10% A at 14.1 min, dwell at 10% A for 14.1-15 min. For reversed phase separation, 5 μL of each sample was injected onto a Waters HSS T3 column (150 x 2.1 mm, 1.8 μm particle size). Column temperature was held at 60°C. Mobile phase consisted of (A) 4:6 (v/v) methanol:water and B 1:9 (v/v) methanol:isopropanol, both containing 0.1% formic acid and 10 mmol/L ammonia. Using a flow rate of 0.4 mL/min, the LC gradient consisted of: Dwell at 100% A at 0 min, ramp to 80% A at 1 min, ramp to 0% A at 16 min, dwell at 0% A for 16-20 min, ramp to 100% A at 20.1 min, dwell at 100% A for 20.1-21 min. MS data were acquired using negative and positive ionization using continuous scanning over the range of m/z 150 to m/z 2000. Data were analysed using an in-house developed lipidomics pipeline written in the R programming language (http://ww.r-project.org), as previously described^56^. All reported lipids were normalized to corresponding internal standards according to lipid class, as well as to freeze-dried tissue weight. Lipid identification has been based on a combination of accurate mass, (relative) retention times, fragmentation spectra, analysis of samples with known metabolic defects, and the injection of relevant standards.

### Proteomics

After the transfer of both solvent layers, the remaining protein pellet was dried under a stream of nitrogen. Proteomics sample preparation was performed using a Thermo Scientific™ EasyPep™ MS Sample Prep Kit (A40006), according to the kit’s instructions. Briefly, 200 µL lysis buffer was added to each sample and a Thermo Scientific™ Pierce™ BCA Protein Assay (23225) was performed according to kit instructions to determine protein content. For each sample, 100 µg of protein was transferred to a new 2 mL tube. Samples were reduced and alkylated at 95°C for ten minutes, followed by a two-hour incubation at 37°C with a Trypsin/Lys-C protease mixture. After sample clean-up with the Peptide Clean-up Plate, samples were dried under nitrogen at 60°C, before being resuspended in a 100 µL mixture of 97:3 (v/v) water:acetonitrile, containing 0.1% formic acid.

Samples were kept at 12°C during analysis and 10 µl of each sample was injected. Injection order for samples was random, with injections of a pooled sample at the start and end, as well as at varying intervals throughout the series. Chromatographic separation was achieved on a Waters™ Acquity UPLC, using a Waters™ Acquity UPLC BEH C18 Column (130Å, 1.7 µm, 2.1 mm X 50 mm) (186002350), equipped with a Waters™ Acquity UPLC Vanguard BEH C18 precolumn (186003975). Column temperature was held at 60°C. Mobile phase consisted of (A) water and (B) acetonitrile, both containing 0.1% formic acid. Using a starting flow rate of 0.5 ml/min, the LC gradient consisted of: Dwell at 3% B for 0-0.1 min; Ramp to 40% B at 4.3 min; Ramp to 80% B at 4.31 min with a flow rate of 0.85 ml/min; Dwell at 80% B for 4.31-4.40 min with a flow rate of 0.85 ml/min; Ramp to 3% B at 4.50 min with a flow rate of 0.6 ml/min; Dwell at 3% B for 4.5-5 min with a flow rate of 0.5 ml/min.

MS data were acquired with a Bruker timsTOF Pro 2 using positive ionization in DIA-PASEF mode as previously reported^57^. A detailed report of the MS method is provided in Supplemental Methods.

DIA-PASEF data files were processed using DIA-NN version 1.8.1^58^, using an empirically generated spectral library for human proteins provided by Bruker Daltonics. The following DIA-NN options were enabled: Reannotate, Contaminants, N-term M excision, C carbamidomethylation, MBR, No shared spectra. Other settings were as follows: Missed cleavages: 1; Peptide length range: 7-30; Precursor charge range: 1-4; Precursor m/z range: 300-1800; Fragment ion m/z range: 200-1800; Precursor FDR (%): 1; Mass accuracy: 0.0; MS1 accuracy: 0.0; Scan window: 0; Protein names (from FASTA); Double-pass mode; Quant UMS (high precision); RT-dependent; IDs, RT & IM profiling; Optimal results. Data were normalized in DIA-NN using MaxLFQ^59^.

### Statistics

Statistical analysis and visualization of the acquired data were done in in the R programming language (http://ww.r-project.org) using the Limma^60^, ggplot2^61^, ropls^62^ and mixOmics^63^ packages. All comparisons between BTHS affected individuals and non-failing donors were performed using a Bayes moderated t-test using the Limma package, after imputation of missing values using a KNN algorithm. Adjusted p-values were subsequently obtained using a Benjamin-Hochberg correction. For other comparisons, for instance within the patient group (i.e. transplant vs autopsy), or comparisons concerning summed analytes (e.g. lipid classes) or grouped analytes (e.g. electron transport chain proteins), methods are described in the text and figures. In general, analytes containing more than 10% missing values were excluded from analysis, unless the missing values were group specific.

## Acknowledgements

We wish to thank all the individuals affected with BTHS, family members and the staff from all departments that contributed to this study. In particular, we would like to express our sincere gratitude to the Barth Syndrome Foundation for their invaluable assistance. This work is financially supported by Human measurement models 2.0: for health research on disease and prevention, from NWO (no. 18953). S.M. was supported by an International Postdoc grant from Independent Research Fund Denmark (1057-00039B).

## Author contributions

Conceptualization: B.V.S., A.S.P., P.J.S., G.S.S., J.vd.V., R.H.H., S.M. Data collection: B.V.S., A.S.P., M.M.T., Y.R.J., D.C., B.J.B., P.M.L.J. Data analysis: B.V.S., A.S.P., M.M.T., P.J.S., M.v.W., I.M.H., C.M.E.L., J.B.v.K., S.R.P., C.R.J., F.M.V. Writing the manuscript: B.V.S., A.S.P. Funding acquisition: J.v.d.V., R.H.H., S.M.

