## Supplemental methods 1 for "Integrated Multi-Omics Mapping of Mitochondrial Dysfunction and Substrate Preference in Barth Syndrome Cardiac Tissue"

### Main

| | | | |
|---|---|---|---|
| **Polarity:** | Positive | **TIMS enable:** | On |
| | | **Scan Mode:** | dia-PASEF |
| **Mass Range from:** | 100 m/z | **Mass Range to:** | 1700 m/z |
| **1/K0 Start:** | 0.60 Vs/cm² | **1/K0 End:** | 1.57 Vs/cm² |
| **Rolling Average:** | On | **Rolling Average No.:** | 10 |
| **View:** | Expert | | |

### Mode - General

| | | | |
|---|---|---|---|
| **Enable TIMS:** | On | **Mark as Calibration Segment:** | Off |
| **Save Spectra:** | Save Frame | **Mass Spectra Peak Detection:** | Use Max. Intensity |
| **Absolute Threshold:** | 10 | **Intensity Threshold:** | Absolute |
| **Intensity Threshold:** | 5000.00 | | |

### Mode - TIMS

| | | | |
|---|---|---|---|
| **ICC:** | Off | **Target:** | 2.0 Mio. |
| **imeX mode:** | Custom | **Resolution:** | Custom |
| **1/K0 Start:** | 0.60 Vs/cm² | **1/K0 End:** | 1.57 Vs/cm² |
| **Ramp Time:** | 100.0 ms | **Spectra Rate:** | n/a Hz |
| **Advanced Parameters:** | On | | |
| **Lock accumul. to mobility range:** | On | **Lock Duty Cycle to 100 %:** | On |
| **Accumulation Time:** | 2.0 ms | **Duty Cycle:** | n/a % |
| **Cycle Time:** | n/a ms | | |

### Source

| | | | |
|---|---|---|---|
| **Source:** | VIP-HESI | | |
| **End Plate Offset:** | 500 V | **Capillary:** | 3500 V |
| **Nebulizer:** | 2.5 Bar | | |
| **Dry Gas:** | 8.0 l/min | **Dry Temp:** | 240 °C |
| **Probe Gas Temp:** | 300 °C | | |
| **Probe Gas:** | 4.0 l/min | **Exhaust:** | On |

### Tune - General

| | | | |
|---|---|---|---|
| **Deflection 1 Delta:** | 70.0 V | | |
| **Funnel 1 RF:** | 300.0 Vpp | **Funnel 2 RF:** | 200.0 Vpp |
| **isCID Energy:** | 0.0 eV | **Multipole RF:** | 500.0 Vpp |
| **Ion Energy:** | 5.0 eV | **Low Mass:** | 200.00 m/z |
| **Collision Energy:** | 10.0 eV | **Collision RF:** | 1500.0 Vpp |
| **Transfer Time:** | 60.0 µs | **Pre Pulse Storage:** | 12.0 µs |
| **High Sensitivity Detection:** | Off | **Stepping:** | Off |

### Tune - Processing

| | |
|---|---|
| **Denoising Mode:** | n/a |

### Tune - TIMS

| | | | |
|---|---|---|---|
| **Dt1 (Defl. Transfer -> Capillary Exit):** | -20.0 V | **Dt2 (Defl. Discard -> Defl. Transfer):** | -160.0 V |
| **Dt3 (Funnel 1 In -> Defl. Transfer):** | 110.0 V | **Dt4 (Accu. Trap -> Funnel 1 In):** | 110.0 V |
| **Dt5 (Accu. Exit -> Accu. Transfer):** | 0.0 V | **Dt6 (Ramp Start -> Accu. Exit):** | 55.0 V |
| **Funnel 1 RF:** | 450.0 Vpp | **Collision Cell In:** | 300.0 V |
